# Mechanisms of dexamethasone-induced bone toxicity in developing bone: a single-cell perspective

**DOI:** 10.1101/2025.01.13.632523

**Authors:** Kelly Warmink, Philip Lijnzaad, Tristan C.W. Vermaat, Lisa Jansen, Marloes C.C. van Gend, Baojie Zhang, Aleksandra Balwierz, Natalie Proost, Marieke van de Ven, Johannes H.M. Merks, Thanasis Margaritis, Claudia Y. Janda

**Author notes:** Lead contact information: Claudia Y. Janda, PhD, Princess Máxima Center for Pediatric Oncology, Heidelberglaan 25, 3584 CS Utrecht, The Netherlands.

## Abstract

Glucocorticoids, such as dexamethasone, are essential for treating severe childhood conditions, including cancer, organ transplantation, and inflammatory disorders. However, their long-term use can impair bone development, posing risks to pediatric bone health, which is vital for lifelong skeletal integrity. A mechanistic insight on how glucocorticoids negatively impact bone could improve decision-making in patient care to improve the quality of life for pediatric cancer patients and survivors. In this study, we aimed to elucidate the molecular mechanisms underlying dexamethasone-induced bone toxicity in developing bones using single-cell transcriptomics. We treated skeletally immature C57BL/6JRj mice with dexamethasone for 28 days, and assessed the bone architecture with micro-computed tomography, and characterized bone and bone marrow cells from the femurs using single-cell RNA sequencing. Our findings revealed a marked reduction in osteoblast and chondrocyte cell populations and impaired function of pre-osteoblasts. Additionally, dexamethasone adversely affected B cell subsets, significantly depleting early B cell progenitors while allowing some further developed immature B cells to persist. These cellular changes were accompanied by reduced longitudinal bone growth, compromised bone architecture, and increased bone fragility at the highest doses of dexamethasone. Interestingly, unlike observations in adults, dexamethasone did not enhance osteoclast activity in our model. Overall, our study suggests that the adverse effects of dexamethasone on bone development are primarily due to its impact on osteoblastic, chondroblastic and B cell lineages, disrupting the critical signaling crosstalk between the cells necessary for bone development and hematopoiesis.

**Layman summary:** Glucocorticoids, like dexamethasone, are vital for treating severe childhood illnesses but can harm bone development when used long-term. This study investigated how dexamethasone affects bone health in young mice. Using advanced techniques, we found that dexamethasone reduced key bone-building cells (osteoblasts and chondrocytes) and weakened their function. It also disrupted immune cell development in the bone marrow, especially early B cells. These changes led to weaker, more fragile bones without increasing bone breakdown, unlike in adults. The findings highlight the need to carefully balance treatment benefits and risks for children to protect their bone health and overall well-being.

## Introduction

Glucocorticoids are widely used for treating various diseases, including cancers, organ transplantation, and autoimmune disorders. These drugs have potent immunosuppressive properties, making them indispensable for managing inflammation and immune responses. In pediatric oncology, glucocorticoids are essential components of acute lymphoblastic leukemia (ALL) and lymphoma therapy and are used to alleviate tumor edema- and therapy-related side effects, such as nausea, allergic reactions, and other immune-related side effects. However, prolonged glucocorticoid use, especially at higher doses as often required in pediatric oncology, is associated with significant side effects, particularly bone-related complications like growth retardation, loss of bone mineral density (BMD), bone fragility, necrosis, and an increased risk of fractures^1,2^. Clinical data show that reduced BMD is common among ALL patients undergoing therapy, with deficits emerging as early as the first month of chemotherapy, even in those patients with normal BMD at diagnosis^3^. Additionally, low BMD often persists post-treatment, and up to every third patient experiences fractures^4,5^. The association of glucocorticoids with bone-related adverse events in ALL patients has been demonstrated in some studies and has recently been summarized by Velentza and colleagues^1^. Treating fractures in these patients is challenging due to the high risk of implantation failure in osteoporotic bone, potentially resulting in chronic pain, limited mobility, and long-term disability. Furthermore, pediatric ALL and lymphoma patients often achieve high survival rates, leading to a long life expectation. This highlights the extended duration of survivorship and the potential long-term effects of treatment.

Dexamethasone and other glucocorticoids exert their effects by binding to the intracellular glucocorticoid receptor expressed in most cell types, including bone (progenitor) cells^6^. The glucocorticoid-receptor complex acts as a transcription factor, activating or repressing the transcription of glucocorticoid-responsive genes. At physiological concentrations, glucocorticoids support bone formation and the accumulation of bone mass by directing the differentiation of mesenchymal stem/stromal cells (MSCs) and stromal progenitors toward the osteoblastic lineage, while inhibiting differentiation into adipocytes^7^. However, when administrated at therapeutic concentrations, glucocorticoids reduce bone formation and increase bone resorption, thereby decreasing overall bone mass and elevating fracture risk^8^. Several mechanisms have been identified by which glucocorticoids directly or indirectly affect skeletal cells, which are comprehensively reviewed by Velentza and Gado and colleagues^1,8^. For example, high concentrations of glucocorticoids inhibit the differentiation of MSCs and osteoblast progenitors towards the osteoblastic lineage, while promoting their differentiation into adipocytes by decreasing the expression of key transcription factors essential for osteoblast differentiation, like runt-related transcription factor 2 (*Runx2*) and Sp7/Osterix (*Sp7*). Furthermore, glucocorticoids inhibit pre-osteoblast maturation by suppressing Wnt and BMP signaling, both of which are crucial for osteoblast differentiation. Additionally, glucocorticoids impair osteoblast function by reducing the expression of bone matrix proteins, including collagen type I and osteocalcin. They also promote apoptosis of osteoblasts and osteocytes through caspase-3 activation and cytochrome C release. Additionally, glucocorticoids stimulate osteoblasts to express RANKL, increasing osteoclast activity and bone resorption. Together, these mechanisms disrupt the intricate balance between bone formation and resorption, resulting in detrimental effects on bone structure, metabolism, and bone function.

The underlying mechanisms of glucocorticoid-induced bone toxicity have predominantly been investigated using cell lines and mouse models involving skeletally mature mice. However, there may be significant differences in the mechanisms, severity, and duration of dexamethasone’s effects between children and adults, as pediatric patients have developing bones characterized by high turnover rates and ongoing remodeling^6,9^. In developing bones, bone growth relies on chondrocyte proliferation and differentiation, followed by endochondral ossification in the epiphyseal plates. Dexamethasone in pediatric patients may disrupt these critical processes of bone growth and remodeling. Yet, the specific molecular mechanisms by which glucocorticoids negatively impact bone formation and resorption in developing bones have not been thoroughly investigated, despite the high need for improved treatment options with fewer side effects in these patients.

Recent advancements in single-cell RNA sequencing (scRNA-seq) have enabled detailed analysis of the cellular composition of adult mouse bone tissue, revealing a diverse array of cell types, including immune cells, MSCs, osteoblasts, chondrocytes, fibroblasts, and various other cellular subtypes^10–12^. In this study, we employed scRNA-seq to investigate the molecular mechanisms underlying dexamethasone-induced bone toxicity in skeletally immature mice, aiming to characterize the effects of dexamethasone on specific cell populations and key bone biomarkers. The insights could enhance patient care and the quality of life for pediatric patients and cancer survivors.

## Results

To investigate the effects of glucocorticoid treatment on developing bones, skeletally immature mice (6-week-old, male, C57BL/6JRj) were injected daily for 28 days with 0 mg/kg (vehicle), 5 mg/kg (DEX 5), 20 mg/kg (DEX 20) or 50 mg/kg (DEX 50) dexamethasone. The age of these mice corresponds approximately to 11.5 years in human terms^13^, which is within the age spectrum for pediatric ALL cases. The 5 mg/kg dose of dexamethasone in mice, approximately equivalent to 0.41 mg/kg in humans, is close to the dose received by pediatric ALL patients during the induction phase (about 0.2-0.3 mg/kg/day depending on the child’s weight). However, the 50 mg/kg dose is significantly higher. After treatment, bone architecture was assessed using micro-computed tomography (μCT), and bone and bone marrow cells were analyzed by scRNA-seq and histology (Fig. 1a). Notably, the DEX 50 group exhibited significant body weight loss during treatment (Fig. S1a), with two mice reaching the humane endpoint, indicating that this dose approaches the maximum tolerated dose for skeletally young C57BL/6JRj mice. Furthermore, there was a trend toward reduced growth in body length among the dexamethasone-treated mice compared to the vehicle group (Fig. S1b). Spleen weight decreased across all dexamethasone-treated groups (Fig. S1c). Although not significant, a similar trend was observed for thymus weight (Fig. S1d), suggesting potential impact on immune cell development and activation. Treatment duration, dosing, and analysis is summarized in Table S1.

**Figure 1:**
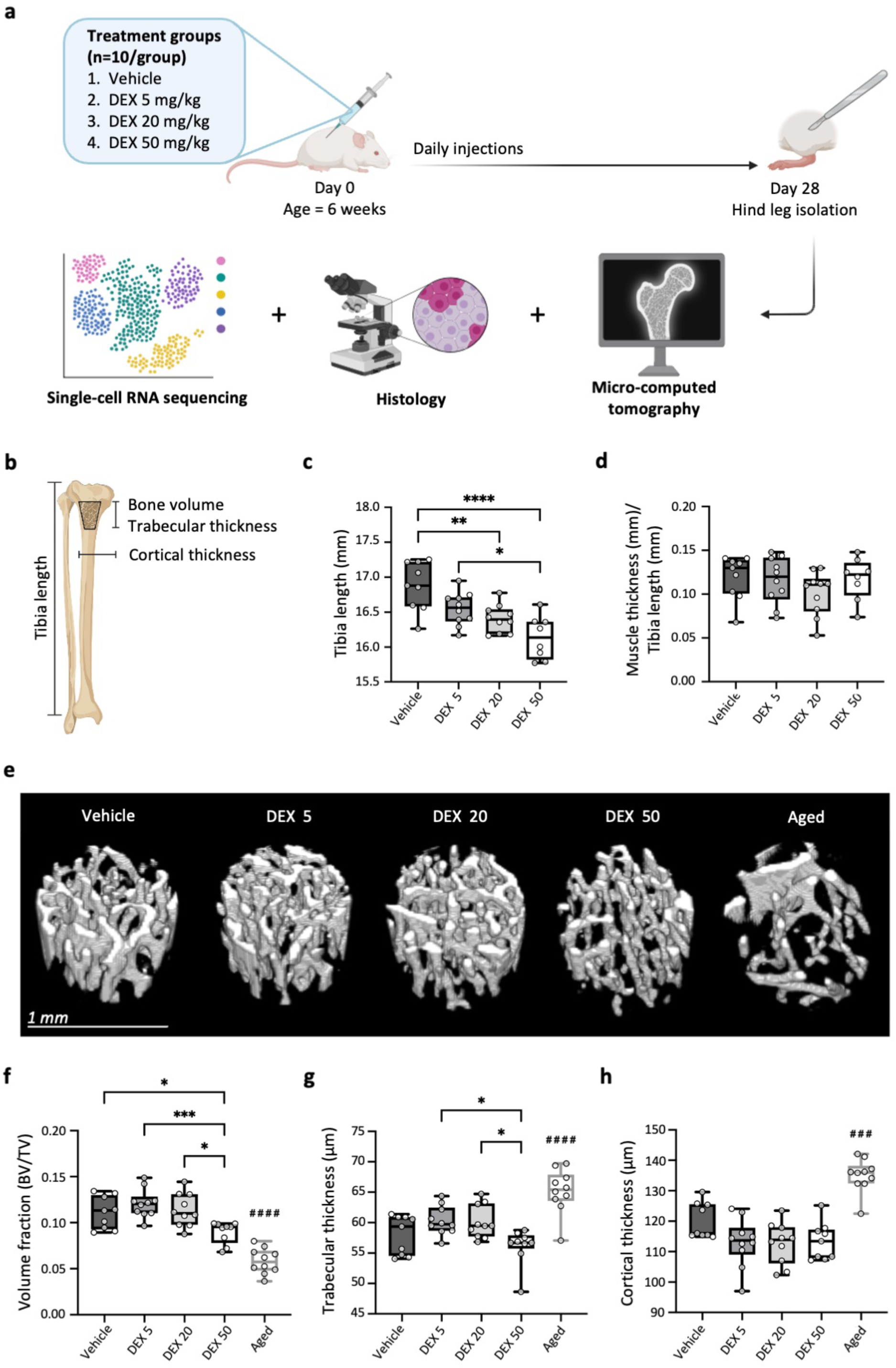
Experimental design and effects of dexamethasone on tibial growth and bone microstructure in skeletally immature mice. **a)** Experimental design illustrating the different treatments of skeletally immature mice with dexamethasone or vehicle. Bone architecture and cellular composition of the bone and bone marrow were analyzed using µCT, histology, and scRNA-seq. **b)** Overview of µCT measurements performed on the mouse tibia. **c)** Box plot depicting tibia length measured by µCT. **d)** Box plot showing muscle thickness quantified as mid-tibia grey area thickness relative to tibia length by µCT. **e)** 3D visualization of the trabecular bone area in the tibia metaphysis used to evaluate bone volume fraction (BV/TV). **f)** Box plot showing bone volume fraction. **g)** Box plot illustrating trabecular thickness. **h)** Box plot showing cortical thickness measured 1.5 mm below the tibia growth plate. Box plots display the median, interquartile range (first to third quartile), and full range (minimum to maximum). Statistical significance is indicated as follows: *p < 0.05, **p < 0.01, ***p < 0.001, ****p < 0.0001. (*) denotes differences between treatment groups, and (#) indicates differences between the aged and vehicle groups.

### Dexamethasone adversely affects longitudinal growth and bone architecture

Bone growth impairment is a well-documented side effect of long-term glucocorticoid therapy in pediatric patients^1,6,9^. To assess the impact of dexamethasone on skeletal development, we examined the tibial growth using µCT. We observed a dose-dependent decrease in total tibia length, with higher dexamethasone concentrations leading to more pronounced reduction (Fig. 1b-c). In contrast, muscle thickness, measured as mid-tibia µCT grey area thickness relative to tibia length, showed no significant differences (Fig. 1d).

Bone volume loss is one of the main bone-related side effects in patients undergoing dexamethasone treatment^1,6,9^. To further explore the effects of dexamethasone on bone volume and architecture, we conducted μCT measurements on the proximal tibia metaphysis (Fig. 1b). We found that the bone volume fraction(BV/TV), and trabecular thickness were significantly and trending lower in DEX 50 mice compared to the vehicle-treated group, respectively (Fig. 1e-g). However, these parameters remained comparable to the vehicle group in the DEX 5 and DEX 20 groups (Fig. 1e-g). Furthermore, the cortical thickness measured 1.5 mm below the growth plate did not significantly vary across the groups (Fig. 1b,h). Given that trabecular bone is more metabolically active than cortical bone, the stronger effect of dexamethasone on the trabeculae was expected. These findings suggest that in young mice, alterations in bone microstructure detectable by μCT occur only at the highest dexamethasone dose of 50 mg/kg, while impaired longitudinal growth is evident even at the lower dose of 5 mg/kg.

To contextualize the dexamethasone-induced bone loss in skeletally young mice, we compared the bone architecture of the proximal tibia metaphysis of dexamethasone-treated mice to that of skeletally old mice (aged; 58-week-old, male, C57BL/6JRj mice from the same breeder), as age is the best-documented risk factor for bone loss across various species, including human and rodents. Compared to dexamethasone-treated and vehicle controls, the BV/TV was significantly lower in aged mice, while trabecular and cortical thickness was significantly higher (Fig. 1e-h). This result suggests distinct underlying molecular mechanisms driving bone loss due to age versus dexamethasone. In summary, our results indicate that dexamethasone treatment negatively impacts both longitudinal bone growth and bone architecture by reducing bone volume fraction and trabecular thickness in skeletally immature mice.

### Dexamethasone-induced bone fragility is unrelated to increased osteoclast activity

Long-term glucocorticoid treatment is known to contribute to bone fragility^1,6,9^. In adults, dexamethasone-induced bone loss is attributed to an imbalance between bone formation and resorption, characterized by reduced osteoblast activity and increased osteoclast activity^14^. Consistently, our study found that dexamethasone treatment increased tibia subchondral plate perforations, characterized by full-depth holes in the normally dense bone layer beneath the cartilage (Fig. 2a-b). The observed impaired bone development was not linked to increased osteoclast activity. This is evident by the tartrate-resistant acid phosphatase (TRAP) staining, which detects osteoblast activity, remaining consistently low in the tibia epiphysis across all treatment groups (Fig. 2c,e). Moreover, TRAP activity in the tibia metaphysis showed a decreasing trend with dexamethasone treatment, although this was not statistically significant (Fig. 2d-e). These findings suggest that bone fragility and impaired bone development seen in these skeletally immature mice are not driven by enhanced osteoclast activity.

**Figure 2:**
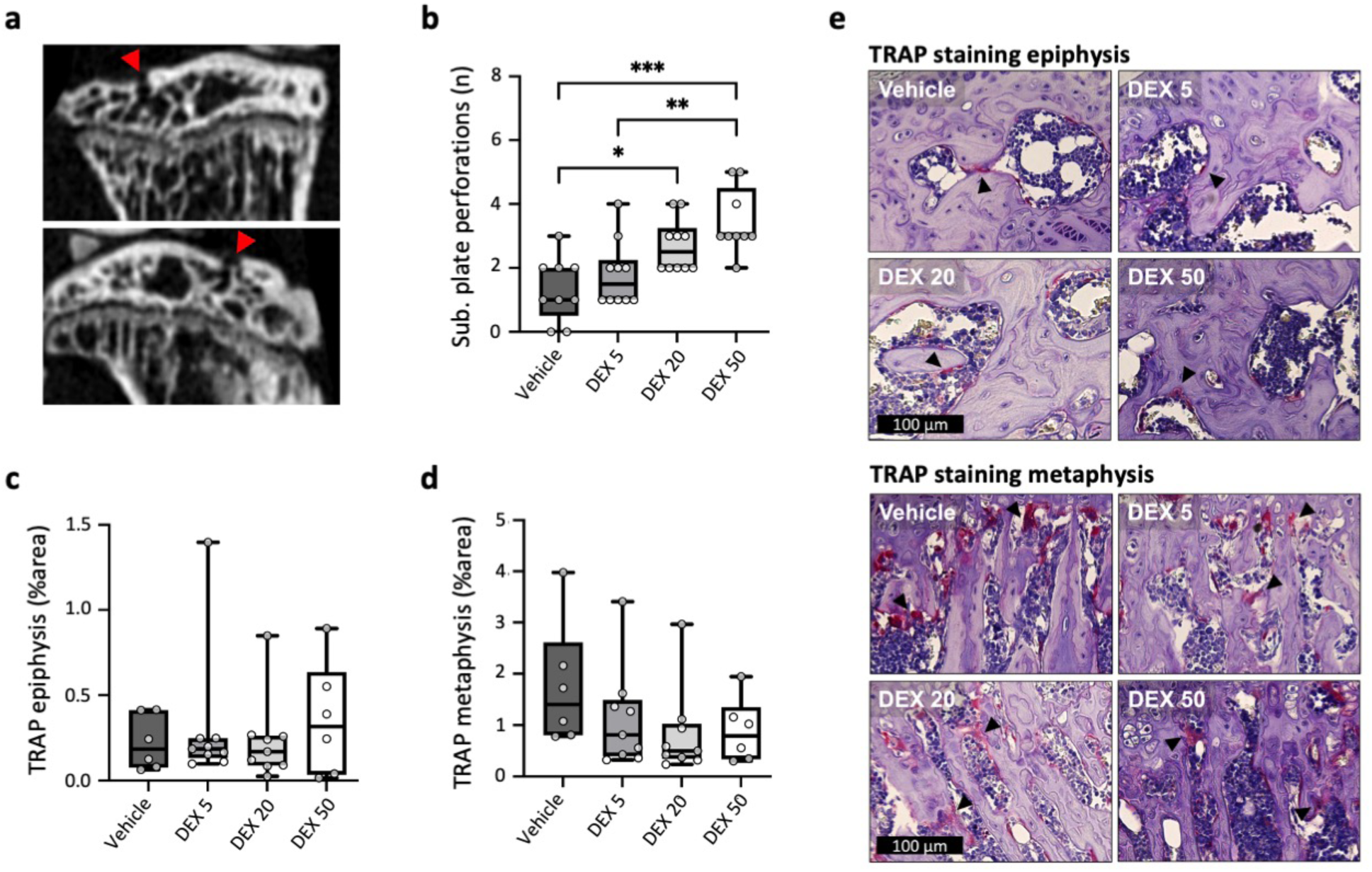
Effects of dexamethasone on bone fragility and osteoclast activity. a) Frontal (top) and sagittal (bottom) sections of µCT scans showing subchondral plate perforations, highlighted by the red arrows. **b)** Quantification of subchondral plate perforations observed in µCT scans across the entire subchondral plate surface. **c-d)** Osteoclast activity, as indicated by TRAP staining, in the c) tibia epiphysis and d) tibia metaphysis, quantified as the percentage positive staining within the total area of interest. Box plots display the median, interquartile range (first to third quartile), and full range (minimum to maximum). Statistical significance is denoted as follows: *p < 0.05, **p < 0.01, ***p < 0.001, ****p < 0.0001. **e)** Representative images of TRAP staining for each treatment group, with positive TRAP staining indicated by black arrows. The images show the epiphysis (top) and metaphysis (bottom).

### scRNA-seq revealed distinct bone and bone marrow cell populations

To deepen our understanding of the molecular mechanisms driving dexamethasone-induced bone toxicity, we performed scRNA-seq on the femurs from dexamethasone-treated and vehicle-treated mice, and a separate group of aged mice (Fig. 3a). After data acquisition, we merged the datasets from the different groups and applied rigorous quality control measures to minimize bias and enhance biological accuracy. The number of transcripts per cell was used as a proxy for cell quality. Low transcript counts indicated poor cell quality, while excessively high counts suggested doublet formation. To ensure data integrity, we set broad cutoffs for transcript counts (≥500, and ≤40,000) and applied filtering on cell barcodes and genes to exclude low-quality cells, including those with >25% mitochondrial transcripts (indicative of dying cells) and >5% hemoglobin transcripts (signifying erythrocyte and erythrocyte precursor contamination). This approach yielded 42,957 cells, unevenly distributed across the three experimental conditions and two control groups (Fig. S2a).We performed cell clustering using the Seurat^15^, focusing on the top 3000 most variably expressed genes. Genes associated with cell cycle phases, hemoglobin, ribosomal proteins, and stress responses were excluded from the analysis. This analysis resulted in distinct cell clusters (Fig. 3b). Immune cell annotations were refined using external bulk-sorted datasets. Non-immune cell annotations were refined using Seurat by projecting the non-immune cell populations onto the mouse bone marrow stroma dataset from Baryawno *et al*.^12^, as described by Stuart *et al.*^16^. Ultimately, we identified 24 distinct cell populations, including 8 stromal cell populations and 16 immune cell populations, which were confirmed through marker gene expression analysis (Fig. 3c, S2b). These populations included bone marrow-resident immune cells, such as neutrophils, B cells, monocytes, and their precursors, as well as non-immune cells like MSC-like cells, osteoblasts, fibroblasts, blood vascular endothelial cells (BEC), and lymphatic endothelial cells (LEC) (Fig. 3b-c). The relative number of cells passing quality control decreased in samples exposed to higher dexamethasone doses. Despite the initial enrichment of CD45^-^ cells to a 1:1 ratio before sequencing, all groups showed increased CD45^+^ cell count (Fig. S2a). Additionally, some clusters displayed mixed cell signatures, like NK/NKT/T and monocyte/DC clusters (Fig. 3b).

**Figure 3:**
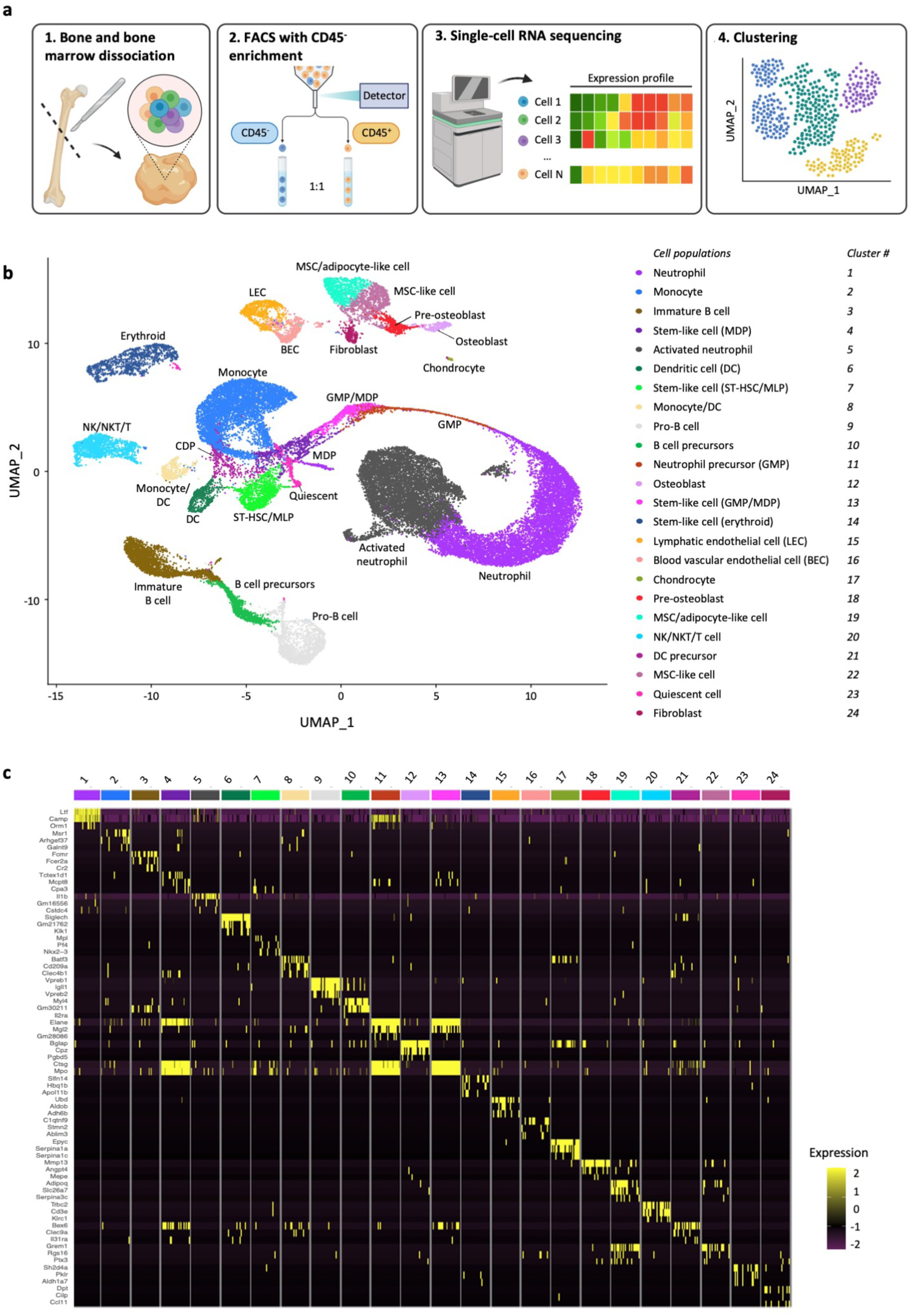
scRNA-seq analysis of femurs from dexamethasone-treated mice. a) Schematic overview of the scRNA-seq experiment design. **b)** UMAP plot showing the final annotation of all identified cell clusters. Clusters with mixed cell signatures are indicated with a forward slash. **c)** Heatmap showing the top 5 most variably expressed genes for each cell cluster.

### Dexamethasone depletes osteoblast cell counts and impairs pre-osteoblast function

To investigate the effects of dexamethasone at the cellular level, we analyzed cell populations using Uniform Manifold Approximation and Projection (UMAP) for cluster analysis, focusing on differences in cell numbers and differential gene expression among treatment groups (Fig. 4a-b, S3a-b). This analysis yielded several key findings.

**Figure 4.**
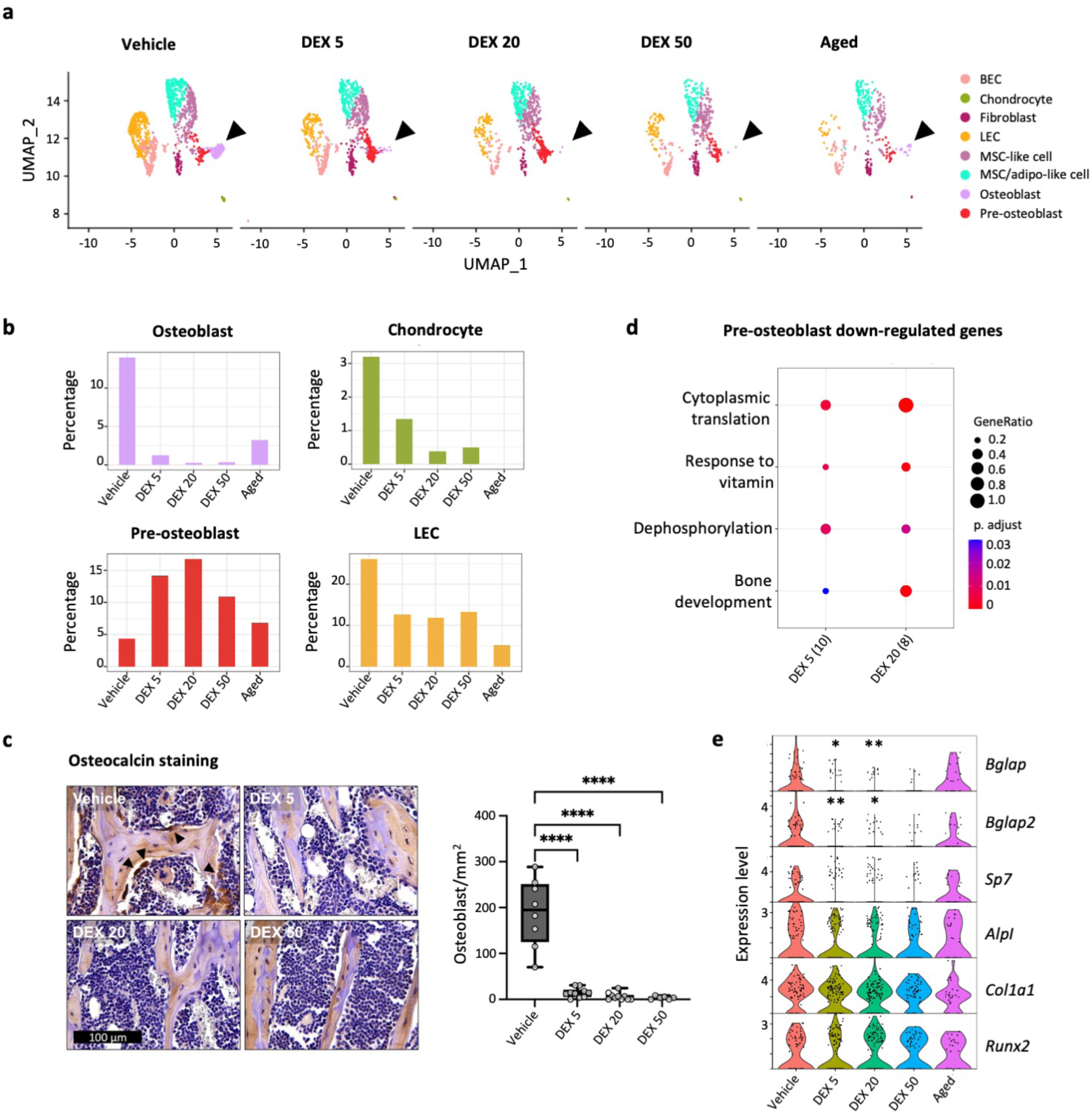
Effects of dexamethasone on endothelial and mesenchymal lineage cells. a) UMAP plot showing all CD45^-^ cells across different treatment group. **b)** Bar graphs showing the percentage of osteoblasts, pre-osteoblasts, chondrocytes and lymphatic endothelial cells (LECs) across different treatment group. **c)** Quantification of osteoblasts per mm^2^ based on osteocalcin immunohistochemistry staining; black arrows indicate osteocalcin-positive osteoblasts lining the bone. **d)** Downregulated Gene Ontology (GO) terms associated with the pre-osteoblast cell cluster. The dot size relates to the number of differentially expressed genes. The total number of differentially expressed genes is indicated in brackets for each treatment condition. **e)** Violin plot showing the expression levels of osteoblast-related genes across treatment groups. Statistical significance is indicated as follows: *p < 0.05, **p < 0.01, ***p < 0.001, ****p < 0.0001.

Firstly, dexamethasone treatment markedly reduced the proportions of both osteoblasts and chondrocytes across all treatment groups compared to vehicle-treated mice (Fig. 4a-b). This reduction was also observed in aged mice. To validate the scRNA-seq results, we performed immunohistochemistry staining for osteocalcin, an abundant non-collagenous bone matrix protein specifically expressed in osteoblasts, to quantify osteoblast cell counts in bone sections. Consistently, we observed a marked reduction in osteoblasts following dexamethasone treatment (Fig. 4c), confirming the scRNA-seq findings.

Interestingly, while scRNA-seq analysis indicated an increase in the percentage of pre-osteoblasts, particularly in the DEX 5 and DEX 20 groups (Fig. 4b), further Gene Ontology (GO) terms analysis revealed a decrease in expression of genes related to bone development and turnover, including processes like dephosphorylation, response to vitamin, and cytoplasmic translation (Fig. 4d, S3c). This suggests that although the relative number of pre-osteoblasts increased, their functional capacity was impaired. For instance, there was a significant downregulation of bone development-related genes *Bglap* and *Bglap2,* which encode osteocalcin, in the DEX 5 and DEX 20 groups (Fig. 4e, S3c). Although a similar decrease was observed in the DEX 50 group, the low number of analyzed pre-osteoblast cells limited statistical analysis. Additionally, markers for osteoblast development, including *Sp7* and Alkaline phosphatase (*Alpl),* also showed a similar trend of downregulation, while genes like Collagen Type I *(Col1a1)* and *Runx2* did not exhibit significant differences between groups within the pre-osteoblast cell cluster (Fig. 4e). Taken together, these findings indicate that dexamethasone treatment leads to a near-complete depletion of mature osteoblasts in skeletally immature mice and significantly impairs the differentiation and function of pre-osteoblasts.

### Dexamethasone decreases lymphatic endothelial cell count and function

Secondly, we observed that dexamethasone treatment decreased the percentage of cells in the LEC cluster (Fig. 4b). This reduction was accompanied by a significant downregulation of genes related to cytoplasmic translation and other GO terms associated with protein production in LECs (Fig. S3d-f). LECs have recently been implicated in bone regeneration, with their function found to decrease in aged animals^17^. These findings suggest that dexamethasone-induced reduction in LEC cell counts could impair healthy bone development in skeletally immature mice.

### Dexamethasone significantly decreases pro-B and immature B cells in the bone marrow

We further interrogated the effects of dexamethasone on immune cells within the bone marrow (Fig. 5a, S4a-b). Overall, the number of immune cells, particularly of B cell lineage cells, in the DEX 50 group was low, prompting us to focus our analysis on the DEX 5 and DEX 20 groups (Fig. S2a).

**Figure 5:**
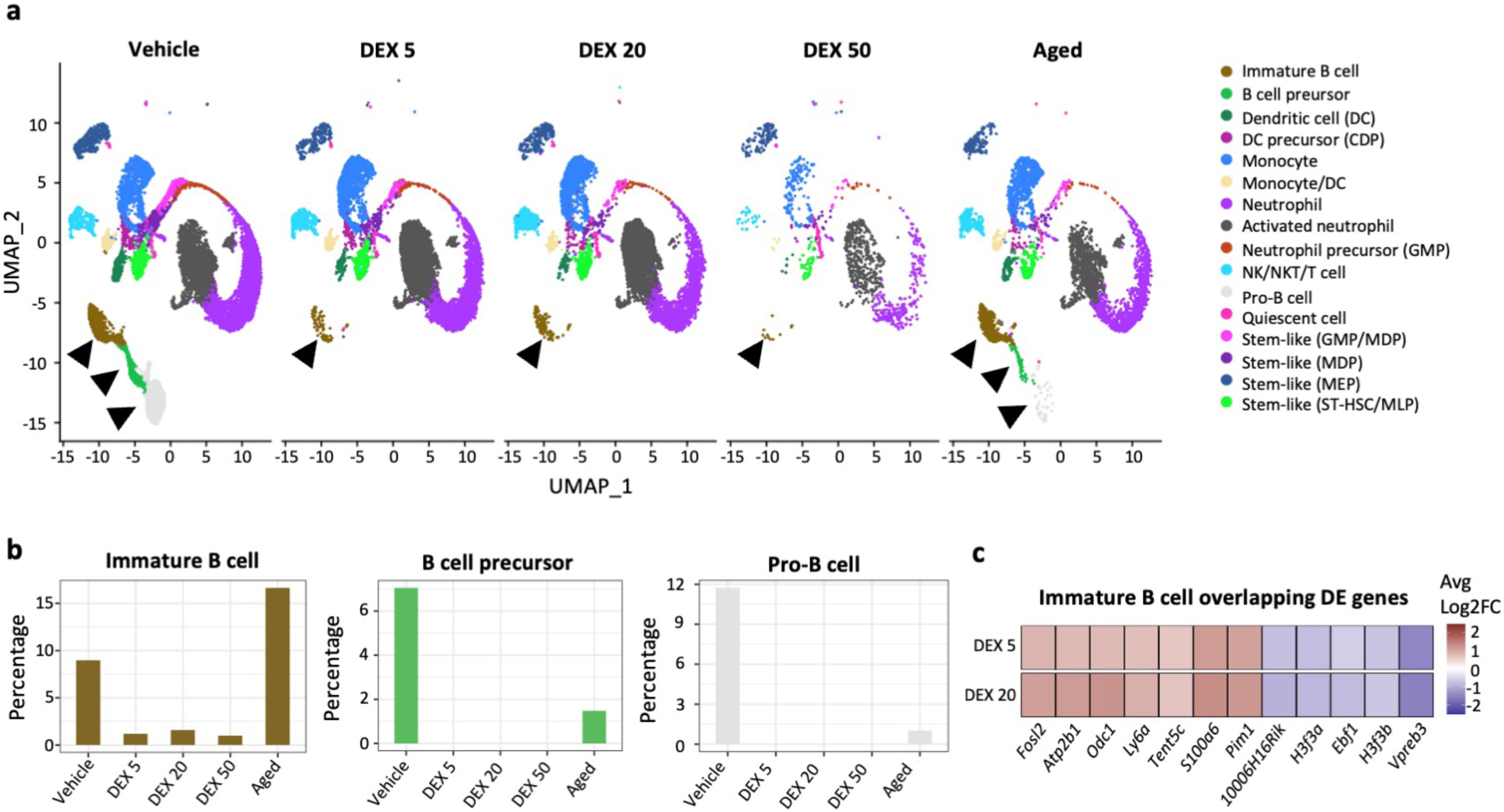
Effects of dexamethasone on the B cell lineage. a) UMAP plot showing all immune cells across treatment groups, with black arrows highlighting B cell clusters. **b)** Proportion of pro-B cells, B cell precursors, and immature B cells across different treatment groups. **c)** Differentially expressed genes in immature B cells in the DEX5 and DEX 20 treatment groups compared to the vehicle group.

Most notably, within the immune cell population, early developing B cells (*i.e.*, the pro-B cell cluster and the B cell precursors cluster) were depleted in all dexamethasone treatment groups while some further developed immature B cells persisted (Fig. 5a-b). As it was not possible to perform differential gene expression analysis on B cell clusters in the dexamethasone-treated groups to confirm cell typing of the different cell clusters, we performed a trajectory analysis on B cell clusters from the vehicle group (Fig. S4c). All B cell clusters expressed the marker gene *Cd19*, and the pro-B cells showed expression of early B cell markers such as *Kit, Spn,* and *Dntt* (Fig. S4d). Genes involved in B cell activation, affinity maturation, and B cell receptor development, including *H2.Eb1, H2.Aa, H2.Ab1, CD74*, *Ighd, Bank1*, and *Cd83* were among the top eight genes upregulated along the pseudotime trajectory (Fig. S4e)^18^. Conversely, genes selectively expressed in pro-B cells, like *Vpreb1* (*CD179a*) and *Igll1* (*CD179b*), were downregulated along the pseudotime trajectory (Fig. S4e).

We then assessed the impact of dexamethasone on gene expression in the immature B cells of the DEX 5 and DEX 20 groups. We observed the significant downregulation of genes essential for maintaining immature B cell commitment and survival, such as *Vpreb3*, *Ebf1*, *H3f3a*, and *H3f3b,* in dexamethasone-treated mice compared to vehicle-treated controls (Fig. 5c)^19–21^. However, genes associated with cell survival, proliferation, and apoptosis, like *Odc1*, *Pim1*, *S100a6*, *Atp2b1*, *Tent5c, Fosl2,* and *Ly6a*, were upregulated in response to dexamethasone. As mature B cells typically leave the bone marrow upon completing their development, their presence is usually minimal compared to immature B cells, which is consistent with the absence of mature B cells in our dataset. In summary, these findings demonstrate that dexamethasone treatment in young animals has a detrimental effect on B cell subsets in the bone marrow, leading to a near depletion of early B cell progenitor cells, while some cells of the more developed immature B cell subset persist.

### Dexamethasone increases bone marrow neutrophil activation and innate immunity

Furthermore, we observed an increase in the percentage of activated neutrophils, which is accompted by an overall increase of neutrophils, following dexamethasone treatment, with the DEX 50 mice having more than double the number of activated neutrophils compared to the control mice (Fig. 6a). Despite the changes in the percentage of the activated neutrophil cluster, the neutrophil cluster remained largely the same (Fig. 6a). Analysis of differentially expressed (DE) genes revealed that both neutrophils and activated neutrophil clusters showed similar gene expression across the DEX groups compared to the control group (Fig. 6b-c). Specifically, dexamethasone led to the upregulation of inflammatory-related genes, including *Cxcl2, Nfkbia, Nfkbiz,* transcription factor *Jun,* and the neutrophil-associated protease *Asprv1* in both neutrophil populations (Fig. 6b-c). On a broader scale, DE genes in neutrophils and activated neutrophils were linked to processes related to cytokine production, chemotaxis (Fig. S5a), and pathogen-associated molecular pattern recognition (Fig. S5a-b). Additionally, the activated neutrophils upregulated genes associated with keratinocyte proliferation, which, in the context of neutrophils, are more likely involved in granule formation. These include Leucine-rich α2 glycoprotein (*Lrg1*), a component of neutrophil granules that get secreted upon activation^22^, and Cathepsin L (*Ctsl*), which plays a role in granule formation^23^ (Fig. S5b). In addition, *Klf9* has been identified as a key regulator of the transcriptomic response to cortisol in zebrafish^24^.

**Figure 6:**
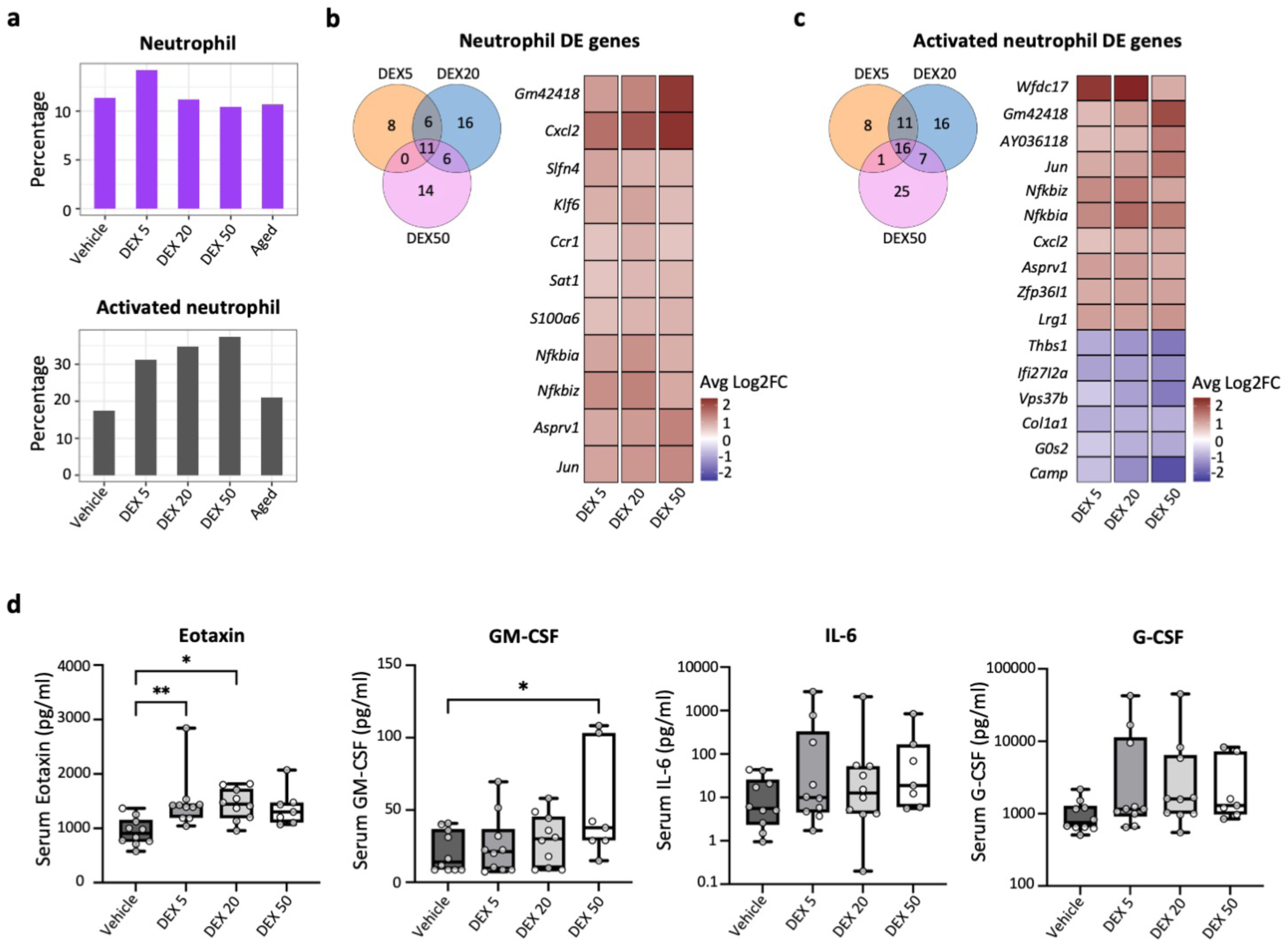
Effects of dexamethasone on neutrophils. a) Percentage of neutrophils and activated neutrophils among all CD45^+^ cells across different treatment groups. **b)** Venn plot summarizing the number of differentially expressed genes between the vehicle and DEX groups in the neutrophil cluster, highlighting the average log-fold change of the 11 genes shared between all dexamethasone-treated mice. **c)** Venn diagram summarizing the number of differentially expressed genes between the vehicle and DEX groups in the neutrophil cluster, highlighting the average log-fold change of the 16 genes shared between all dexamethasone-treated mice. **d)** Endpoint serum cytokine concentrations of eotaxin, GM-CSF, IL-6, and G-CSF across treatment groups. Box plots show the median, interquartile range (first to third quartile), and the full range (minimum to maximum). Statistical significance indicated as follows: *p < 0.05, **p < 0.01, ***p < 0.001, ****p < 0.0001

In accordance with increased neutrophil activation and upregulated inflammation, the endpoint serum of dexamethasone-treated mice showed elevated levels of systemic Eotaxin and GM-GSF, and the inflammatory cytokines IL-6 and G-CSF showed a similar upward trend, but this was not statistically significant (Fig. 6d). Altogether, these findings suggest that prolonged daily exposure to dexamethasone enhances bone marrow neutrophil activation and stimulates inflammatory cytokine responses in skeletally young mice.

## Discussion

Glucocorticoids cause bone-related side effects in pediatric patients but remain essential for treating ALL and lymphoma^1,6,9^. Understanding the molecular mechanisms driving glucocorticoid-induced bone toxicity during skeletal development could aid in managing growth suppression, bone loss, necrosis, and fracture risk during and after treatment and facilitate decision-making in patient care to improve the quality of life for pediatric cancer patients and survivors. In this study, we investigate the molecular mechanisms underlying dexamethasone-induced bone toxicity using skeletally immature mice. It should be noted that glucocorticoid-induced bone loss manifests in rodents and humans with comparable cellular and molecular characteristics, making mice a suitable first model to study the effects of glucocorticoids using single-cell omics approaches^25^. Through scRNA-seq, we investigated the effects of dexamethasone (5, 20, and 50 mg/kg) on various cell populations within the long bones. Our results reveal consistent cellular alterations across the different doses, including the reduction in osteoblasts, chondrocytes, LECs, and B cell precursors, and an increased activation of neutrophils. These changes were associated with reduced longitudinal bone growth, adverse impacts on bone architecture, and increased bone fragility. Notably, while cellular changes were evident at a daily dose of 5 mg/kg, structural changes in trabecular bone were only detected at the higher daily dose of 50 mg/kg, indicating that cellular alterations precede visible structural changes. Overall, our findings suggest that dexamethasone-induced bone toxicity in young mice involves widespread alterations in the bone and bone marrow microenvironment, affecting osteoblasts, immune cells, and endothelial cells, leading to impaired bone development.

In adults, dexamethasone-induced bone loss is attributed to an imbalance between bone formation and resorption, characterized by reduced osteoblast activity and increased osteoclast activity^14^. In our study with skeletally immature mice, dexamethasone strongly impacted bone-forming osteoblasts, while its adverse effects on bone health were unrelated to osteoclast activity. We observed that dexamethasone significantly reduced osteoblastic lineage cells and downregulated key bone development genes, particularly osteocalcin (*Bglap* and *Bglap2)*. This aligns with clinical observation where therapeutic doses of glucocorticoids rapidly decreased bone formation markers, like osteocalcin, alkaline phosphatase (ALP) and procollagen type I C-terminal telopeptide, secreted by osteoblasts^26,27^. Similarly, in rodents, glucocorticoid exposure decreases bone formation markers, osteoblast numbers, and bone mineralization^28–31^. Intriguingly, glucocorticoids downregulate osteocalcin through the binding of the glucocorticoid-receptor complex to negative response elements in the enhancer region of the osteocalcin promotor^32^, explaining their stronger impact on osteocalcin compared to other bone turnover markers. While BMD measurements are recommended for evaluating glucocorticoid effects on bone, bone turnover markers are also being explored^33^. Given that glucocorticoids directly, yet differentially, affect the expression of the bone turnover markers, a thorough understanding of the underlying mechanisms is crucial for accurate data interpretation.

Unlike in mature bone, dexamethasone treatment in our skeletally immature mice did not alter osteoblast-related TRAP activity, suggesting that in developing bones, the detrimental effects of dexamethasone are primarily due to reduced bone formation rather than increased bone resorption. Due to challenges in isolating pre-osteoclasts and osteoclasts, we could not compare their RNA profiles across treatment and age groups to confirm this. Nevertheless, this finding has potential implications for pediatric patients with dexamethasone-induced bone loss, as treatments with osteoclast-inhibiting bisphosphonates primarily target basal osteoclast activity, rather than the increased osteoclast activity. Indeed, bisphosphonate efficacy in pediatric glucocorticoid-induced osteoporosis is variable and generally less potent than in adults^34,35^. However, a placebo-controlled study with 217 participants (4-18 years) with chronic inflammatory rheumatic disease, showed a positive effect of bisphosphonates on bone density, comparable to adults^36,37^. Nevertheless, a Cochrane Systemic Review concluded insufficient evident to recommend bisphosphonates as standard therapy in children^38^. Factors like inflammatory status, age, and bone development stage likely contribute to these variable outcomes and warrant further investigation.

Furthermore, our study revealed that dexamethasone depletes B cell-committed precursors (*i.e*., pro-B cells, B cell precursors, and immature B cells), which are essential for maintaining a functional immune system. Similar effects have been reported in adult mice and humans treated with single or multiple injections of corticosteroids, where the underlying mechanism is thought to involve induced apoptosis across all B cell developmental subsets^39,40^. This apoptotic response is exploited in treating childhood cancers like B cell ALL, where glucocorticoids support remission when combined with chemotherapy^41^. In our study, dexamethasone downregulated key genes involved in B cell commitment and survival pathways (*Ebf1*, *H3f3a*, *H3f3b*), potentially shifting toward T cells differentiation, as suggested by *Ebf1* downregulation and *Ly6a* upregulation. Upregulated genes in immature B cells (*e.g*., *Odc1, Tent5c*) were associated with cell survival and proliferation, suggesting that these genes might help prevent glucocorticoid-induced B cell apoptosis. This could contribute to a shift toward T cell differentiation in the surviving B cell subset. It is well recognized that immune cells within the bone marrow play a crucial role in regulating bone homeostasis and regeneration^42^. Stromal cell populations, many of which originate from mesenchymal lineages, interact with immune cells to regulate the development and function of hematopoietic cells. Among the key cell types within the bone marrow niche that support B cell development are osteoblast progenitors and mature osteoblasts. These cells provide essential signals through direct cell-cell interactions and the secretion of critical cytokines and growth factors, including CXCL12, IL-7, CD147, and Galectin-1 (Gal-1). Additionally, RANKL-RANK signaling is crucial for both osteoclast differentiation and B cell development. In the bone marrow, osteoclasts and B cell precursors express the receptor RANK, while RANKL is produced by osteoblast precursors, mature osteoblasts and other stromal cells. Disruption of this signaling pathway has been shown to lead to reduced B cell populations^43,44^. The decrease in osteoblast populations observed in our study may have contributed to the negative effects of dexamethasone on B cells, and vice versa. These findings underscore the importance of considering the interplay between immune and bone cells in understanding dexamethasone-induced bone toxicity and overall bone health.

## Materials and methods

### Animals

Male 6-week-old C57BL/6JRj mice (n=40, Janvier Labs) were randomized at baseline into four groups (n=10 per group): vehicle, DEX 5, DEX 20, and DEX 50 (Centrafarm B.V., Breda, The Netherlands) in 10ul per gram bodyweight. Mice received daily intraperitoneal injections for 28 days. Body weight was measured daily, and the tail and body length were measured weekly. After four weeks, mice were euthanized, spleens and thymuses were isolated and weighed, and hindlimbs were kept in medium at 4°C for processing. The left hindlimb was cut at the distal femur, cleaned, and fixated in 4% buffered formaldehyde for 24 hours for subsequent μCT imaging and histology. The proximal left femur and the complete right femur were cleaned, dissected, frozen viably and stored at -180°C for scRNA-seq. Furthermore, ten untreated 58-week-old male C57BL/6JRj mice (Janvier Labs) were included for μCT and scRNA-seq analysis.

Mice were housed in individually ventilated IVC cages with maximal five animals per cage, provided with bedding, nesting material, red domes, and *ad libitum* access to water and RM3 chow. Housing conditions included temperature control (21±0.5°C), humidity control (45±2%), and a 12:12 light-dark cycle. Humane endpoints included ≥20% body weight loss, 15% loss within 2 days, circulation or breathing issues, or aberrant behavior/movement. Animals were monitored daily. The study protocol was approved by the ethical committee and animal welfare body under CCD number: AVD3010020198564 Appendix 1 Protocol ID 21.1.10472 and 21.1.10191, complying with European Community guidelines. The protocol was not (pre)registered in an online preclinical study registry.

### Excluded animals

One mouse of the DEX 50 group died unexpectedly after 8 days, likely due to internal bleeding observed in the intestine, and was excluded due to insufficient treatment duration. Two additional DEX 50 mice reached the humane endpoint of 20% body weight loss after 22 and 23 days, and were sacrificed. While tissues were harvested and processed as per protocol, these mice were excluded from measurements potentially affected by the delayed timing (serum, sequencing, and histology) but not μCT measurements. Following consultation with animal caretakers, technicians, and the animal welfare body, the remaining six DEX 50 mice were sacrificed to prevent further suffering and ensure proper harvesting. Consequently, the DEX 50 group in this study includes 9 mice treated for 21-26 days, instead of the intended 28 days. Additionally, one vehicle-group mouse was excluded from μCT and histology due to a bony outgrowth in the proximal tibia. An summary of all mice, treatment, and inclusion per analysis is provided in Supplementary Table 1.

### Micro-computed tomography

The left hindlimbs were imaged using the Quantum FX μCT (PerkinElmer; SAP160036a) at 90kV and 180μA. Two scans were performed: one with a 5mm field of view (FOV) for proximal tibia imaging (3 minutes, isotropic voxel size 9.8μm) and another with a 20mm FOV for distal femur to foot imaging (4.5 minutes, isotropic voxel size 39.1μm). CT images were analyzed in ImageJ (1.47v) using the BoneJ plugin^45^.

The scans with a FOV of 5mm were used to measure trabecular bone thickness (µm) and bone volume fraction. The bone was segmented using a local threshold algorithm (Bernsen algorithm, radius 5). The trabecular bone volume fraction (BV/TV) was calculated by the ratio of trabecular bone volume (BV, mm^3^) and endocortical tissue volume (TV, mm^3^). Regions of interest (ROI) were drawn in transverse plane 103 slides (±1 mm), starting below the growth plate going distal on the tibia by drawing an oval, without reaching the cortical bone, at the first and last image slide and interpolating the ROIs in between. The mean thickness of the trabeculae was measured using the same ROI. The thickness of the cortical bone was determined by measuring the mean thickness of 11 slides (±100µm) in the transverse plane 1.5mm below the tibia growth plate. In addition, the number of perforations in the tibia subchondral plate was counted.

The scans with a FOV of 20mm were used to measure the tibia length and muscle thickness. The scan was rotated to the sagittal plane in a view that showed both the medial malleolus and medial meniscus. The tibia length was measured from the medial meniscus to the medial malleolus. The muscle thickness was defined as the perpendicular line until the edge of the grey area, at half of the tibia length. All μCT analyses were done blinded and in randomized order by 2 independent observers.

### Histology

After μCT imaging, left hindlimbs were processed for histology. Bones were decalcified in 0.5 M EDTA (pH 7.0) for three weeks, refixed weekly in 4% buffered formaldehyde, dehydrated in ethanol (70-100% ethanol), cleared in xylene, and paraffin-embedded in sagittal orientation for 4μm sections. Haematoxylin and eosin (H&E) staining was performed at the Princess Máxima Center.

TRAP staining for osteoclasts involved incubation of the slides for 4 hours at 37 °C, with 2.5 mg/ml Naphtol AS-MX phosphate and 5.5 mg/ml Fast red TR salt in 0.2 M acetate buffer (pH 5.0) with 50 mM tartaric acid. Osteocalcin immunohistochemistry was used to detect osteoblasts. Slides were blocked for endogenous peroxidase using 0.3% H_2_O_2_ (10min, RT) and antigen retrieval was performed in Tris-EDTA buffer (pH 9.0, 30min, 37°C). Primary antibody incubation was overnight (4°C) with 0.1 µg/ml osteocalcin polyclonal antibody (PA5-78870, Invitrogen) or 0.1 µg/ml IgG isotype control (08-6199, Invitrogen). Slides were incubated with anti-rabbit IgG HRP (ab205718, Abcam) (30min, RT) and developed using DAB+ (Agilent, Santa Clara, USA) for 5min. Sections were counterstained with hematoxylin, dehydrated, cleared, and coverslipped.

Images were captured using a Leica DMi8 light microscope, three 20x images were taken per section and analyzed in ImageJ (v2.9.0). TRAP-stained areas were quantified by threshold, and osteocalcin-positive osteoblasts directly neighboring bone tissue were counted. All image acquisition and analysis were done blinded and randomized.

### Serum measurements

On day 28 tail vein blood was collected in serum tubes, centrifuged (1800 RCF, 10min) and the serum snap-frozen in liquid nitrogen. Samples were stored at -180°C until analysis. Serum cytokines and chemokines were measured using a 32-multiplex assay (Merck, MCYMAG-70K-PX32) at the MultiPlex Core Facility, University Medical Center Utrecht, the Netherlands. Analytes with >25% out-of-range values (IL-3, IL-7, IL12p70, LIF) were excluded. IFNγ was below, and LIX above the detection limit. Remaining out-of-range values (6% of all measurements, all below detection limits) were set at ½ of the lower quantification limit.

### Statistics imaging and serum data

Statistical analysis was performed using Prism (v9.3.1, GraphPad) and R (v4.2.1). Differences between treatment groups were assessed with one-way ANOVA followed by Tukey’s multiple comparisons test for μCT, histology, and serum data. Data distribution was checked using Q-Q plots of residuals. Longitudinal data (body mass and length) were analyzed with a mixed-effects model (REML), with time point and treatment group as fixed effects and individual animals as random effect. P-values ≤0.05 were considered significant. Data are presented as boxplots (median, interquartile range, min-max) with significance indicated as *(p≤0.05), **(p≤0.01), ***(p≤0.001), and NS (not significant). Group allocation was blinded during post-life measurements, and all procedures were randomized.

### scRNA-seq sample preparation

Femurs from mice were pooled per treatment group to ensure sufficient cells for scRNA-seq. To enhance downstream reliability (e.g., differential gene expression) femurs from 20 vehicle-treated mice from 2 studies with an identical study protocol were included. Bone fragments were digested in two 30-minutes rounds at 37°C using digestion buffer containing Optimem (Gibco, 12634010), 100 U/mL DNase (Stemcell, 7469), 10 mM HEPES buffer (Gibco, 15630056), 1 mM CaCl_2_, 10 μM ROCK (Y-27632, Biogems, 1293823), and 0.1 mg/mL Liberase DL in the first digestion round (Sigma, 5466202001) or 0.1 mg/mL Liberase TM in the second digestion round (Sigma, 05401119001). Bone fragments were strained (70 μm), washed extensively, and centrifuged (300g, 5min) after each step. Collected cells were resuspended in 5 ml FACS buffer (magnesium-free PBS, 1% BSA, 1 mM EDTA, 10 μM ROCK) and kept on ice. Red blood cell were lysed with RBC buffer (Invitrogen, 4min, RT), washed, and blocked 8 μg anti-CD16/CD32 antibody (rat anti-mouse BD, 553079) (15min). Cells were incubated with 4 μg CD45-FITC (Stemcell, 60161.2, Clone 2.4G2) (30min, 4°C), washed and resuspended in 1 mL FACS buffer. Sorting was performed using a Sony SH800 Cell Sorter (100 μm nozzle chip), with droplet calibration (Sony beads, LE-B3001). Live/death cells were identified via DR (Biolegend, 5μM) and DAPI (Biolegend, 2.5μM) staining. Gates were set using unstained controls for DR+, DAPI-, single cells, CD45^+^ and CD45- populations. CD45^+^ and CD45^-^ cells were sorted and combined in a 50:50 ratio, with 45 000 cells per condition used for scRNA-seq. Cells were processed using the 10x Genomics Chrominum Single Cell 3’ Kit. Libraries were prepared using the 10x Genomics Chromium 3’ Gene Expression solution v3.1 and sequenced on a NovaSeq6000 (Illumina).

### scRNA-seq data processing

Reads were mapped to mouse mm10-2020-A transcriptome and feature count tables were generated using the CellRanger count function. The read10Xlibs function from the SCutils package (v1.110) was used to return a raw Seurat object in R (v4.2.1). Counts for mitochondrial genes are read to determine the mitochondrial percentage, and the percentage of hemoglobin complex gene expression was determined to estimate erythroid contamination. Cells with <500 unique transcripts, >40 000 unique transcripts, >25% mitochondrial transcript percentage, or >5% hemoglobin transcript percentage. Doublets were annotated using scDblFinder (v1.10) on the RNA assay and data slots of the Seurat object^46^. scDblFinder was run separately per library using *nfeatures* = 1 000 and *iter* = 4. Subsequently, cells with a doublet probability above 0.5 were removed. The resulting object was normalized using the SCTransform method^47^ and analyzed using the Seurat R package, version 4.1.1^15^, and the top 3 000 most variably expressed genes were defined. The cell cycle phase of each cell was determined using the CellCycleScoring function in Seurat, using the built-in gene lists. Cell cycle phase correlated genes were determined using metadataCorrelations and derivedCellcycleGenes from the SCutils package. Both cell cycle phase and cell cycle correlated genes were removed from the list of variable genes to avoid biases in cell clustering. Hemoglobin, ribosomal, and stress-related genes were removed from the variable features, resulting in 2799 variable features. The top 35 principal components were used to project in UMAP, using default settings. Clustering was performed using the FindNeighbors and FindClusters functions, with 35 principal components and a resolution of 1.8.

The resulting 46 clusters were annotated using SingleR 1.10.0^48^, with the MouseRNAseqData and ImmGenData reference datasets. The top 5 most variable genes per cluster were used to check and refine the cell annotations. The non-immune cell clusters were confirmed with cell projection^16^, using the mouse bone marrow stroma data by Baryawno et al.^12^.

### scRNA-seq data analysis

Differentially expressed genes were identified per cluster and condition using the FindMarkers function in Seurat, with logfc.threshold = log2(1.5) and min.cells.group = 10. DE genes were filtered on a p.val.adj < 0.1. Top 25 DE genes underwent GO enrichment analysis (Biological Process) with enrichGO from clusterProfiler (v4.8.2)^49^. B-cell trajectory inference was performed by first re-clustering the B-cells separately with a Gaussian mixture model using mclust (v6.0.0) (https://mclust-org.github.io/mclust/reference/Mclust.html). Secondly, Slingshot (v2.8.0) was used to infer the pseudotime^50^. Genes (anti)correlating maximally with the B cell Trajectory pseudotime were identified using Pearson correlation and subjected to gene set over-representation analysis.

### Data availability

Single-cell RNA-seq data have been deposited at GEO with the accession number is GSE281311 and will be made publicly available as of the date of publication. GEO includes astq data, the count tables and the final Seurat object in the form of an RDS file. Any additional information required to reanalyze the data reported is available from the lead contact upon request.

### Declaration of Generative AI usage

In the final steps of the preparation of this manuscript the authors used Grammarly, ChatGPT to improve the language. After using these tools, the authors reviewed and edited the content as needed and take full responsibility for the content.

## Acknowledgements

The presented work is supported by the Foundation Children Cancer Free (KiKa grant 384, C.Y.J.). We thank the Janda group, the Single Cell Genomics Facility, and the van den Heuvel-Eibrink group of the Princess Máxima Center for helpful discussion and feedback, the MultiPlex Core Facility of the University Medical Center Utrecht for performing the multiplex assay, the Princess Máxima Center Flow Cytometry and Imaging Facilities for their assistance.

## Author contributions

C.Y.J. conceived and designed research. N.P. and M.v.d.V. performed and supervised mouse experiments. K.W., L.J., M.C.C.v.G., and B.Z. performed mouse sample processing. K.W. and M.C.C.v.G performed μCT measurements and analysis. K.W. and M.C.C.v.G performed and analyzed immunohistochemistry staining. K.W. and A.B. prepared samples for scRNA-seq. K.W, P.L., T.C.W.V and T.M. analyzed scRNA-seq data. K.W. and C.Y.J. interpreted results and wrote the manuscript. C.Y.J. and J.H.M.M. provided supervision. All authors reviewed and approved the final manuscript.

---

**Supplementary figure 1: Systemic effects of dexamethasone.** a) Changes in body mass and **b)** body length relative to baseline measurements during the 28-day treatment period (error bars represent standard deviation). **c)** Spleen weight measured at the study endpoint. **d)** Thymus weight measured at the study endpoint. Box plots show the median, interquartile range (first to third quartile), and the full range (minimum to maximum). Statistical significance is indicated as follows: *p < 0.05, **p < 0.01, ***p < 0.001, ****p < 0.0001.

**Supplementary figure 2: Final cell counts and marker genes used in scRNA-seq analysis. a)** Absolute cell counts representing the final number of cells that passed quality control, shown for each treatment group. **b)** Marker gene expression profiles to validate and confirm the annotation of cell clusters.

**Supplementary figure 3: Effects of dexamethasone on mesenchymal and endothelial cell lineages. a)** Proportional representation of mesenchymal lineage and endothelial cell types within total CD45- cells across different treatment groups. **b)** Bar graphs showing the percentage of MSC-like cells, MSC/adipocyte-like cells, fibroblasts, and blood vascular endothelial cells (BECs) within the mesenchymal and endothelial lineage across different treatment group. **c)** Netplot showing downregulated Gene Ontology (GO) terms and differentially expressed genes between the vehicle and DEX groups within the pre-osteoblast cell cluster. **d)** Downregulated GO terms identified in the LEC cluster. The dot size relates to the number of differentially expressed genes. The total number of differentially expressed genes is indicated in brackets for each treatment condition. **e)** Netplot displaying differentially expressed genes in the LEC cluster alongside their corresponding downregulated GO terms. **f)** Violin plot showing the module score for genes involved in cytoplasmic translation within lymphatic endothelial cells across treatment groups.

**Supplementary figure 4: Effects of dexamethasone on the B cell lineage.** a) Proportions of various immune cell types across the different treatment groups. **b)** Bar graphs showing the proportion of monocytes, dentritic cells (DC), monocyte/DC, DC precursors, NK/NKT/T cells, quiescent cells and neutrophil precursors across different treatment groups among all CD45^+^ cells. **c)** Pseudotime trajectory of B cell lineage clusters, mapping the developmental progression of B cells. **d)** UMAP plot illustrating the expression of *Cd19, Kit, Spn and Dntt* within B cell clusters. **e)** Heatmap showing the expression of the top 8 genes with the strongest positive and negative correlation with pseudotime.

**Supplementary figure 5: Effect of dexamethasone on neutrophils. a)** Netplot visualizing differentially expressed genes in neutrophils from dexamethasone-treated mice compared to the vehicle group, along with their associated GO terms. **b)** Netplot visualizing differentially expressed genes in activated neutrophils from dexamethasone-treated mice compared to the vehicle group, along with their associated GO terms.

**Table S1**: Summary of the treatment duration, dosing, and analysis inclusion for each mouse.

