## Supplementary figures for "Mechanisms of dexamethasone-induced bone toxicity in developing bone: a single-cell perspective"

### Supplementary figure 1

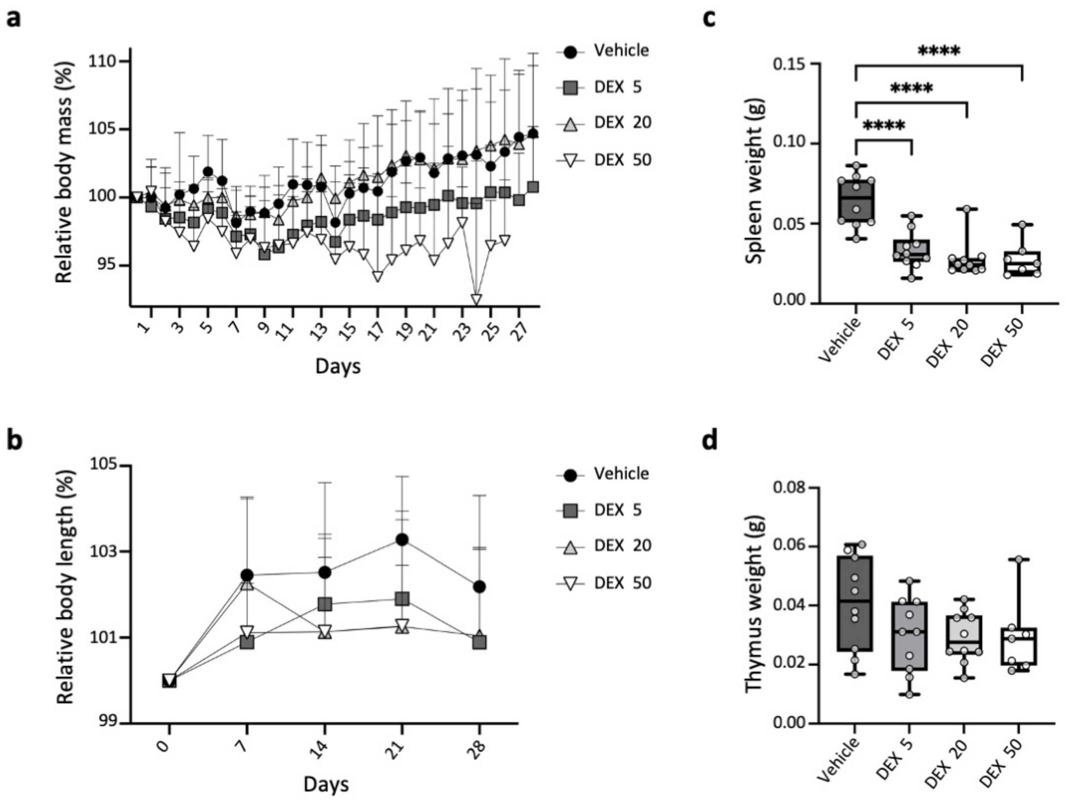

### Supplementary figure 2

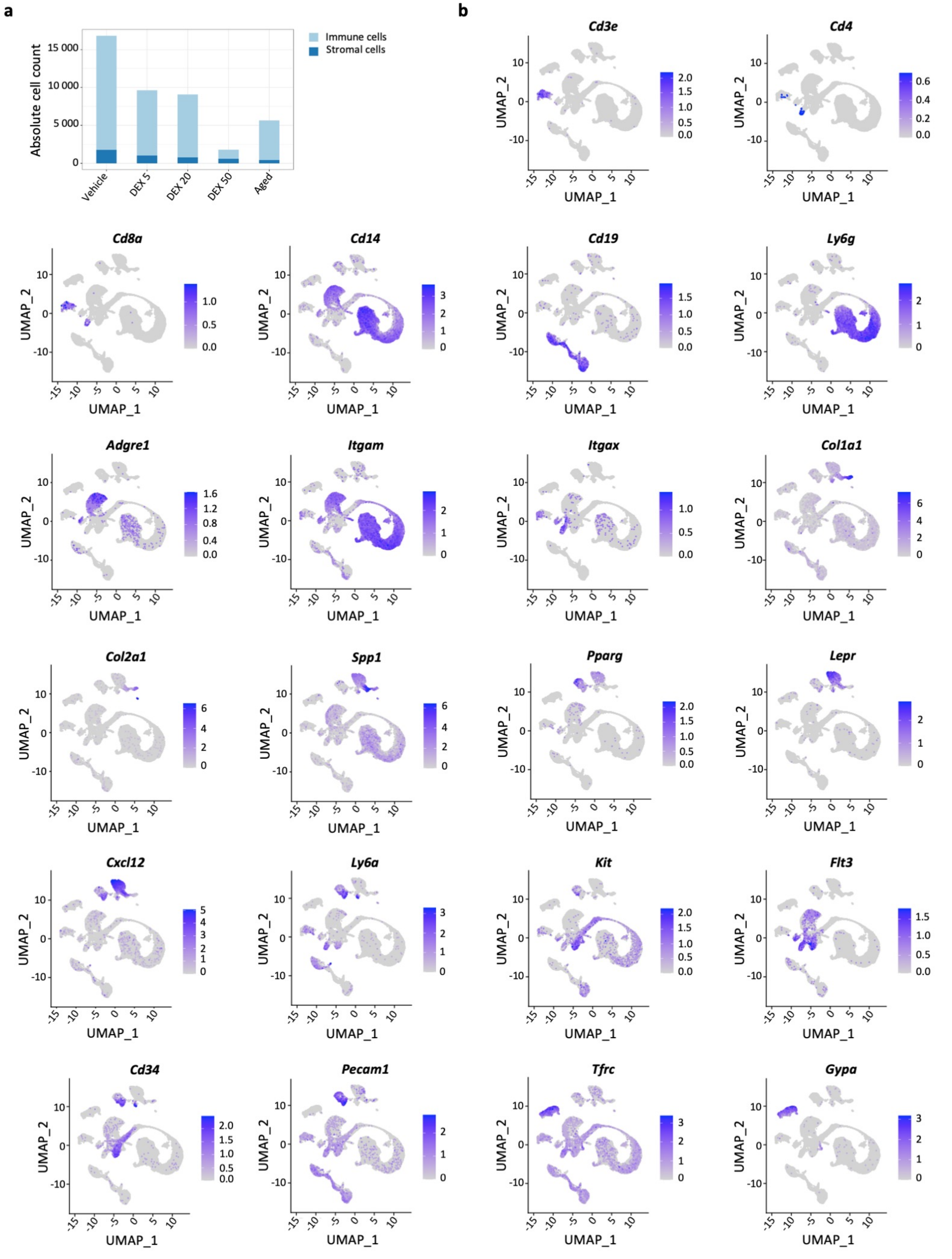

### Supplementary figure 3

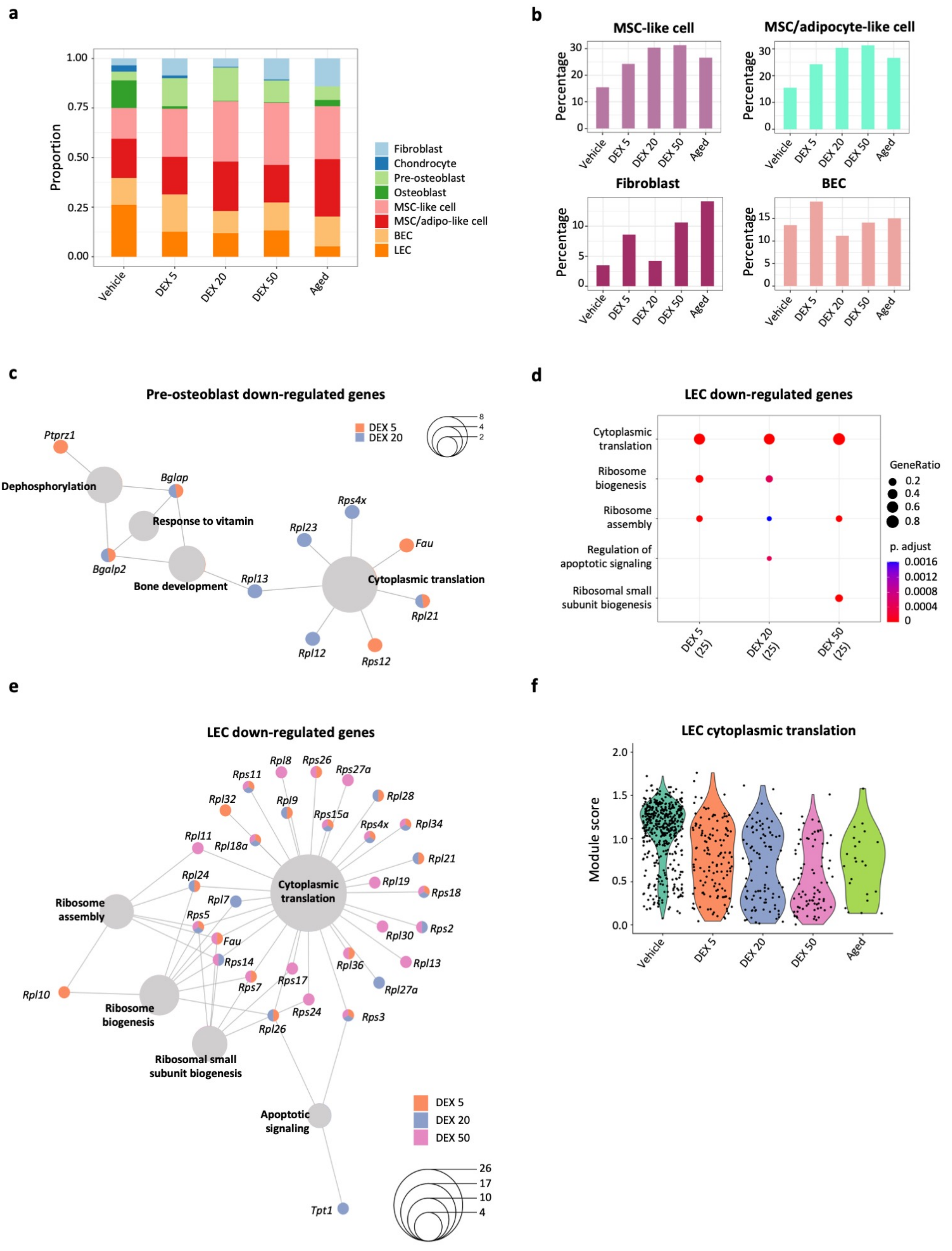

### Supplementary figure 4

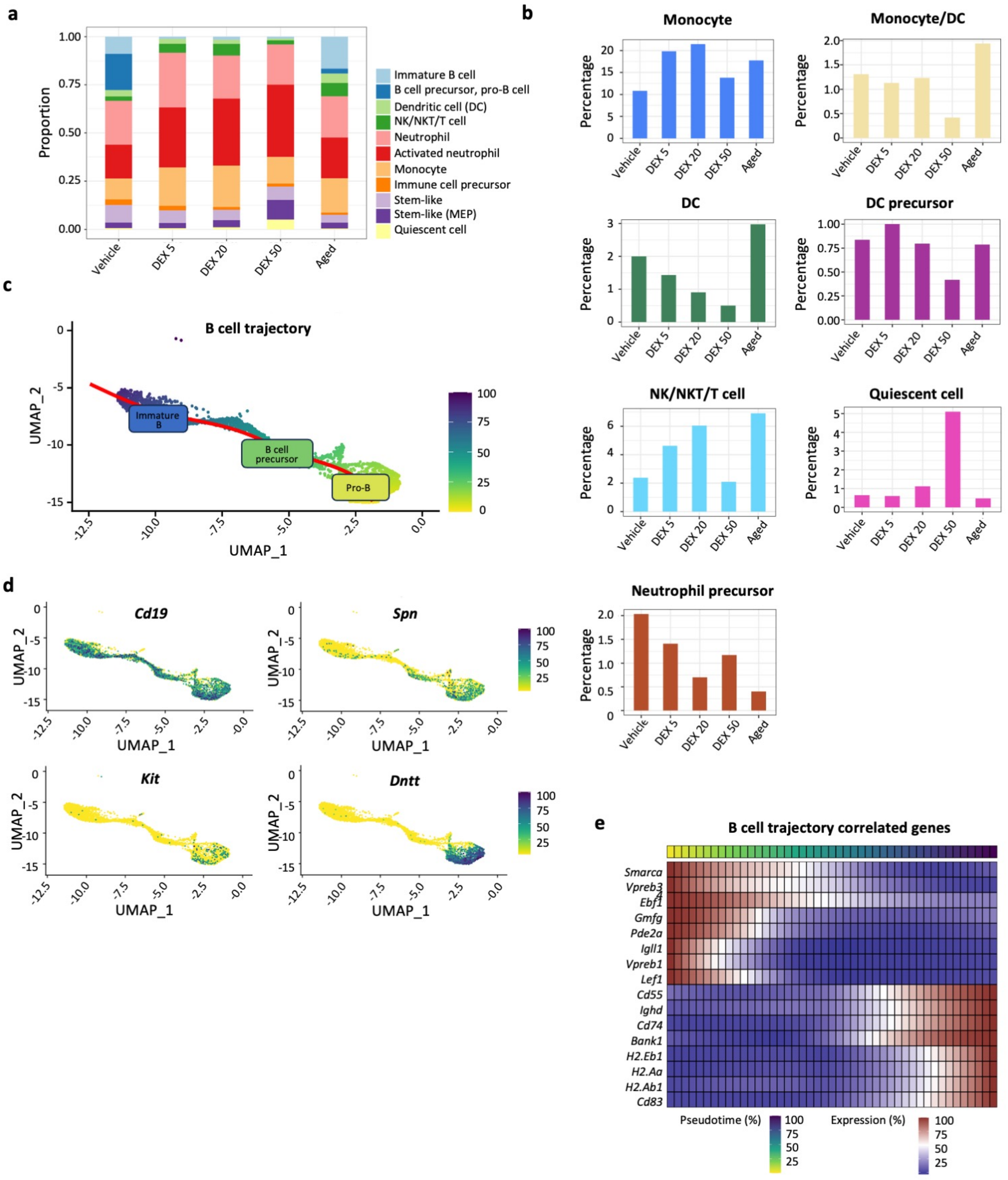

### Supplementary figure 5

**a** Enriched processes associated with upregulated genes in neutrophils in DEX-receiving mice compared to control mice

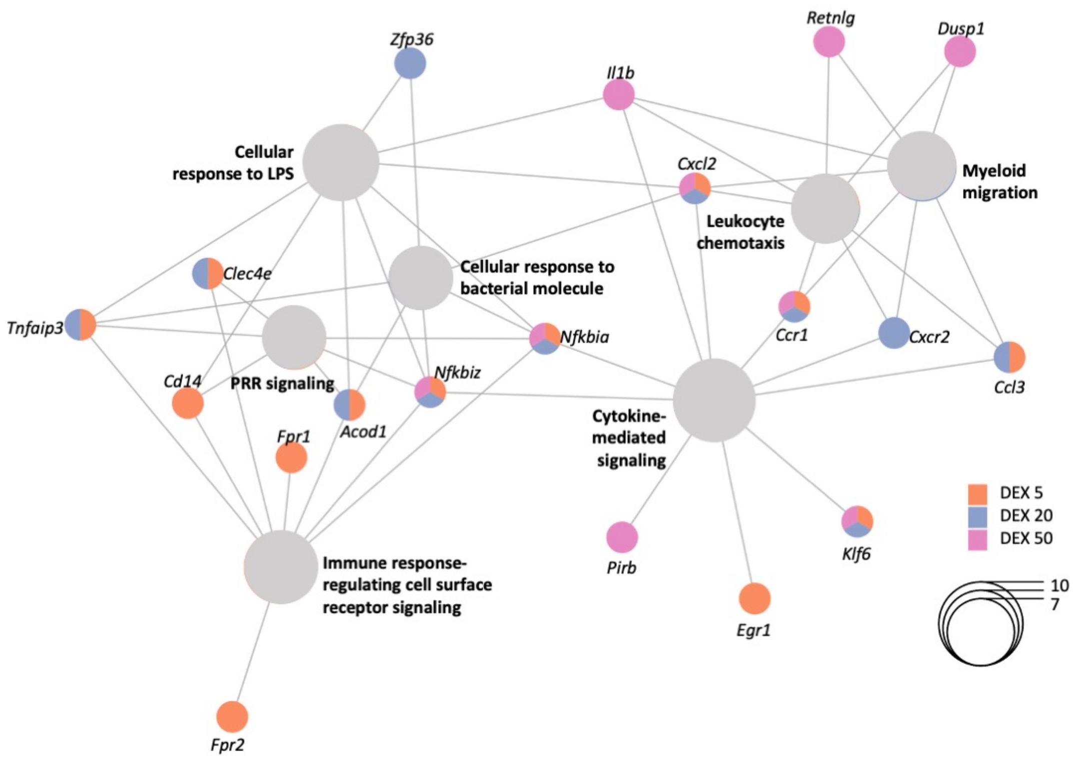


**b** Enriched processes associated with upregulated genes in activated neutrophils in DEX-receiving mice compared to control mice

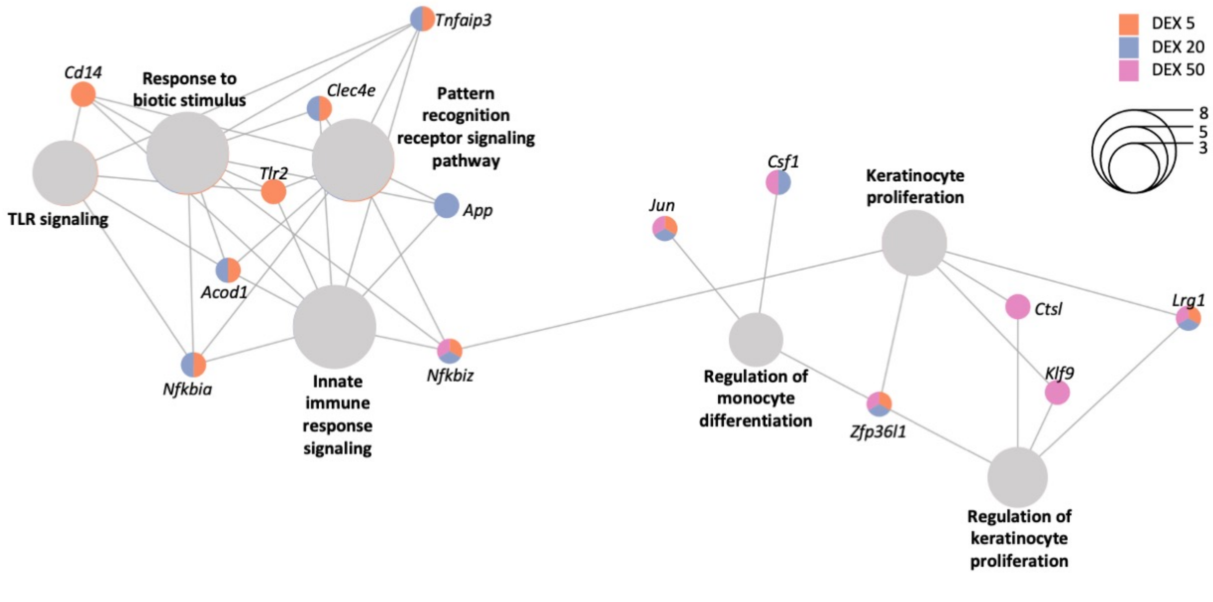

#### Table S1

| Group | Mice | Study time (d) | Reason of death | Body mass and length *Suppl. Fig. 1* | Organ weight *Suppl. Fig. 1* | CT scan (FOV20) *Fig. 1c-d* | CT scan (FOV5) *Fig. 1f-h, Fig 2b* | TRAP *Fig. 2c-d* | scRNAseq *Fig. 3-6* | Osteocalcin *Fig. 4c* | Serum Multiplex *Fig. 6d* |
|---|---|---|---|---|---|---|---|---|---|---|---|
| | | | | **Total animals included per analysis** | | | | | | | |
| Vehicle | n=10 | 28 | endpoint sacrifice | 10 | 10 | 9 (c) | 9 (c) | 8 (c,d) | 10 | 8 (c,d) | 10 |
| DEX 5 | n=10 | 28 | endpoint sacrifice | 10 | 10 | 10 | 10 | 9 (d) | 10 | 10 | 10 |
| DEX 20 | n=10 | 28 | endpoint sacrifice | 10 | 10 | 10 | 10 | 9 (d) | 10 | 10 | 10 |
| DEX 50 | n=1 | 8 | unexpected death (a) | 9 (a) | 7 (a,b) | 8 (a,e) | 9 (a) | 6 (a,b,d) | 7 (a,b) | 6 (a,b,d) | 7 (a,b) |
| | n=1 | 22 | humane endpoint (b) | | | | | | | | |
| | n=1 | 23 | humane endpoint (b) | | | | | | | | |
| | n=4 | 21 | precautionary sacrifice | | | | | | | | |
| | n=3 | 26 | precautionary sacrifice | | | | | | | | |

**Reason exclusion (n excluded)**

a) Excluded from study due to insufficient treatment duration (n=1)
b) Excluded from selected analyses, due to prolonged harvesting duration (n=2)
c) Bone outgrowth visible (n=1)
d) Low section quality (n=1)
e) Faulty scan (n=1)
